# A Multi-Breed Reference Panel and Additional Rare Variation Maximizes Imputation Accuracy in Cattle

**DOI:** 10.1101/517144

**Authors:** Troy N. Rowan, Jesse L. Hoff, Tamar E. Crum, Jeremy F. Taylor, Robert D. Schnabel, Jared E. Decker

## Abstract

**Background:** The use of array-based SNP genotyping in the beef and dairy industries has produced an astounding amount of medium-to-low density genomic data in the last decade. While low-density assays work exceptionally well in the context of genomic prediction, they are less useful in mapping and causal variant discovery. This project focuses on maximizing imputation accuracies to the marker set of two high-density research assays, the Illumina Bovine HD, and the GGP-F250 which contains a large proportion of rare and potentially functional variants (~850,000 total SNPs). This 850K SNP set is well-suited for both imputation to sequence-level genotypes and direct downstream analysis.

**Results:** We find that a large multi-breed composite imputation reference comprised of 36,131 samples with either HD and/or F250 genotypes significantly increases imputation accuracy compared to a standard within-breed reference panel, particularly at low minor allele frequencies. Imputation accuracies were maximized when an individual’s ancestry was adequately represented in the composite reference, particularly with complete 850K genotypes. The addition of rare content from the F250 to our composite reference panel significantly increased the imputation accuracy of rare variants found exclusively on the HD. Additionally, we identify 50,000 variants as an ideal starting density for 850K imputation.

**Conclusion:** Using high-density genotypes on all available individuals in a multi-breed reference panel maximizes imputation accuracy for all cattle populations. Admixed breeds or those sparsely represented in the composite reference are still imputed at high accuracy which will increase further as the reference panel grows. We expect that the addition of rare variation from the F250 will increase the accuracy of imputation at the sequence level.

## Background

High-density single nucleotide polymorphism (SNP) genotyping has driven incredible genetic progress in livestock populations [1–3]. To further increase the predictive abilities of these tests, functional variant discovery has become increasingly important. While many large-effect or Mendelian loci controlling important phenotypes in cattle have been discovered [4–8], moderate and small effect quantitative trait nucleotides (QTN) and other causal variants have proven more challenging to pinpoint. Early genome-wide association studies (GWAS) aimed at identifying these loci were often forced to choose between resolution (number of SNPs) and statistical power (number of individuals). Imputation, the use of statistical models and a reference set of haplotypes to infer the state of missing genotypes, allows studies to genotype large numbers of individuals at low-density and impute to high-density or even full-sequence genotypes [9–11].

Low-to-medium density SNP assays are widely used in genetic evaluations for both beef and dairy cattle. Since the debut of the BovineSNP50 BeadChip (Illumina, San Diego, CA) [12] in 2008 and the BovineHD (Illumina, San Diego, CA) in 2009, more than 3 million dairy cattle have been genotyped in the United States alone on assays derived from these two initial assays [13]. Decker (2015) underlined the value of these commercially-generated datasets for uses outside of genetic prediction [14]. While lower-density assays work exceptionally well in a genomic prediction context, they have a lower resolution for QTL or causal variant detection. Imputation allows these datasets of unprecedented size to be utilized to their full potential. Seabury et al. (2017) found that 50K genotypes provided insufficient resolution for QTL detection, but an analysis using 777K imputed genotypes yielded 14 clear QTL signals [15]. Using these large commercial datasets imputed to high-resolution marker densities will unlock a new level of biological discovery and potentially increase prediction accuracies in cattle [16–19].

To utilize these large datasets to their full potential, maximizing the accuracy of imputation is critical. The most effective imputation software packages for cattle [20] are typically developed for use in human studies aimed at imputing from a high-density genotype panel to full-sequence. As a result, using these programs for imputing directly from low-density to full-sequence, even in cattle breeds with high levels of LD, is ineffective. A “two-step” imputation strategy, first from a low-density assay to a high-density assay and then from imputed high-density to sequence was more effective than a “one-round” imputation from low-density to full-sequence in both cattle and humans [21,22]. Here, we concern ourselves with the first half of the “two-step” imputation process as it can be used for further imputation to sequence, or as an endpoint for downstream analysis. Regardless of its use, maximizing the accuracy of high-density genotype imputation is central to the success of either application.

Initially, SNP assays for cattle were designed with common, evenly-spaced markers that would presumably be in linkage disequilibrium with causal variants [12]. While these assays have performed exceptionally well in genomic prediction schemes, interest in including rare variation into predictions has become increasingly common [9,17,19,23]. Imputation accuracy has been shown to decline rapidly at low minor allele frequencies (MAF), so increasing confidence in the imputation of rare variants has become a priority. Additionally, the majority of imputation optimization studies have focused on the imputation of purebred animals using closely related individuals of the same breed. As large numbers of genotypes for unpedigreed crossbred animals become available, we have to re-evaluate our strategies for imputation in these datasets.

This work focuses on maximizing imputation from various low-density assays to a common set of high-density variants (850K), many of which are rare and potentially functional. We test the effectiveness of a large, multi-breed composite reference for imputation in multiple different beef and dairy cattle populations. We use both well-established and novel measures of imputation accuracy to categorize precisely where, why, and how imputation errors are made. These metrics provide insights for interpretation and situations in which to exercise caution when using imputed variants. Additionally, we explore how starting chip density impacts the accuracy of 850K imputation. Finally, we introduce the GGP-F250 functional genotyping assay as a tool not only for genotyping numerous functional variants but also for increasing the imputation accuracy of rare variants.

## Materials & Methods

To identify the best practices for high-density, chip-level imputation accuracy, we compare the impact of a number of parameters on imputing to the union of two popular research assays; the Illumina BovineHD (Illumina, San Diego, CA), and the GeneSeek Genomic Profiler F250 (GeneSeek. Lincoln, NE) referred to as the HD and F250 respectively. The HD assay contains 777,962 evenly spaced variants at relatively high minor allele frequencies. The F250 assay contains 227,234 variants; 31,392 of which are common variants used for imputation, and another 195,842 potentially functional variants, many of which are rare. These rare alleles make the minor allele frequency distribution of the F250 appear more similar to that of the site frequency spectrum of the bovine genome (**Figure 1**). We used 2,719 animals genotyped on both the F250 and the HD assays, 25,772 animals genotyped on the F250, and 7,218 genotyped on the HD assay.

**Figure 1.**
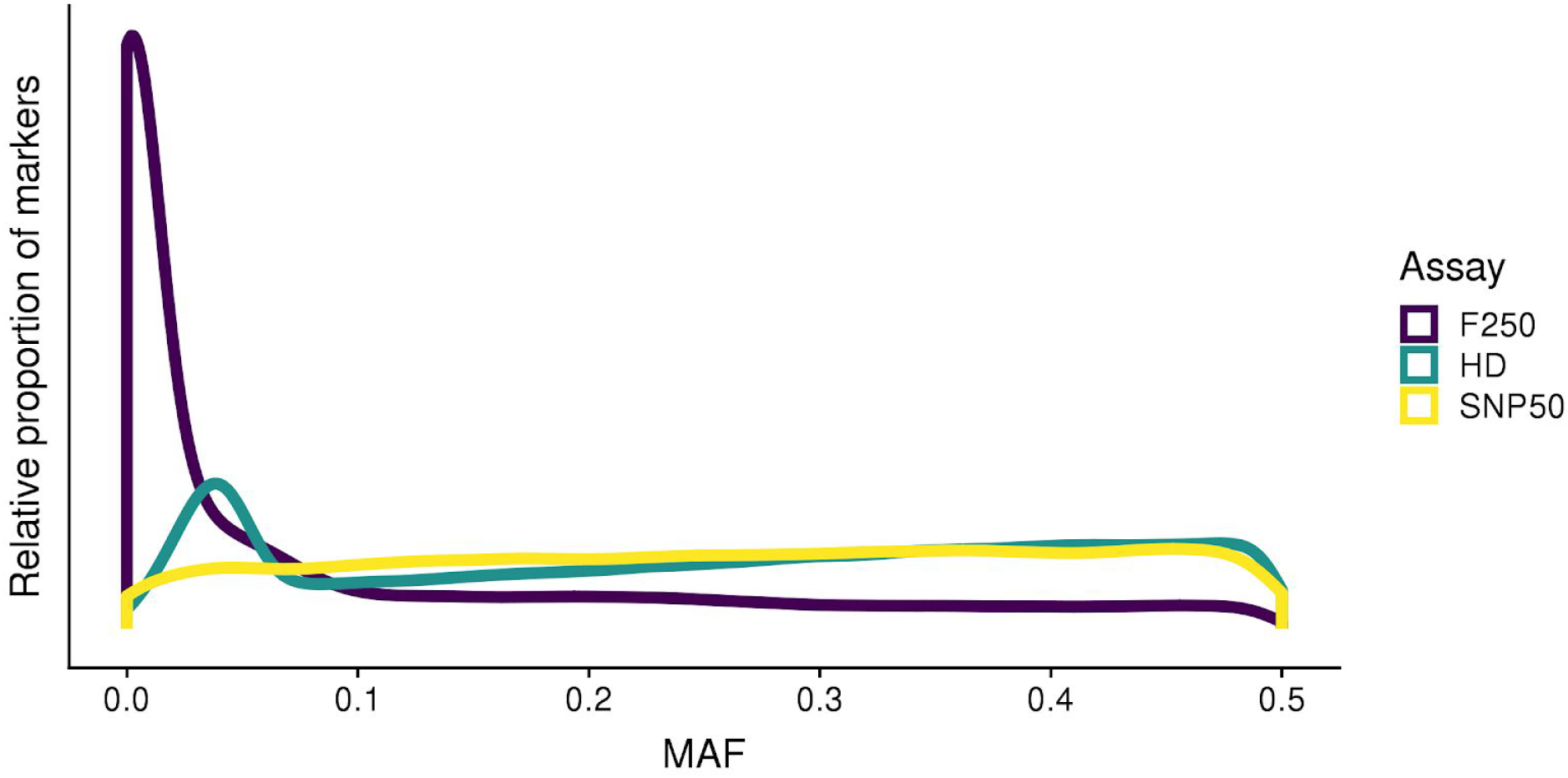
Site frequency spectra for various assay densities. Density plot of minor allele frequencies of SNP50 (yellow), F250 (purple), and HD (blue).

### Quality Control and Filtering

PLINK1.9 [24] was used to filter problem variants and individuals. The SNP positions were based on the ARS-UCD1.2 Bovine reference genome assembly [25]. Non-autosomal variants were removed from the data. Variants and individuals with call rates < 0.90 were removed from testing and reference datasets. Because of the rare nature of many F250 variants, no MAF filter was applied to this data. Due to the divergent nature of this multi-breed dataset, no Hardy-Weinberg Equilibrium filter was applied. PLINK was used to calculate minor allele frequencies at this point for all downstream analyses. Two animals were dropped due to low call rates. The number of variants remaining after filtering for each of the assays tested is shown in **Table 2**.

### Downsampling

To test imputation accuracy, PLINK1.9 [24] was used to downsample 308 animals with observed HD and F250 genotypes to the density of multiple common commercial genotyping arrays; the BovineSNP50 (Illumina, San Diego, CA) [12], GGP-LD, GGP-90KT, GGP-HDv3, and GGP-ULD (all from GeneSeek, Lincoln, NE) and then impute to the union of two high-density research chips (~850K SNPs). All commercial assays tested are almost complete subsets of markers genotyped on the HD (**Supplementary Table 1**).

A maximum of 50 individuals per breed, with complete genotypes on both HD and F250, were chosen to be downsampled to commercial chip densities for imputation testing (**Table 1**). All test set individuals had breed-composition estimates based on the CRUMBLER pipeline [26]. To avoid depleting the reference panel of underrepresented breeds, no more than 50% of a breed’s F250 or HD animals were removed for testing. The remainder of HD and F250 genotypes were used in the composite reference panel (**Table 2**). Due to the uneven representation of certain breeds in the testing dataset, we created three separate datasets for testing different aspects of our imputation pipeline. The first, ALL, uses all 308 downsampled individuals that passed genotype filtering. Because some of the indicine breeds used in our testing dataset are not adequately represented in the imputation reference, or their testing numbers are not large enough to draw conclusions from, we created a dataset of only *Bos taurus* animals, referred to as TAUR, that contained 281 individuals from Angus, Gelbvieh, Hereford, Holstein, Limousin, and Simmental. Finally, we use a dataset of 49 Gelbvieh individuals, referred to as GEL, to compare the accuracy of a breed-specific imputation reference to our composite reference.

**Table 1.** Breed representation of testing set and the proportion of the breed’s total F250 and HD content that they represent.

| **Breed** | **No. for testing** | **%Both** | **%HD** | **%F250** |
| --- | --- | --- | --- | --- |
| Angus | 50 | 27.47% | 2.36% | 0.34% |
| Holstein | 50 | 2.52% | 1.55% | 2.51% |
| Hereford | 44 | 100.00% | 7.18% | 2.34% |
| Simmental | 50 | 42.74% | 10.48% | 2.76% |
| Nelore | 4 | 57.14% | 0.47% | 50.00% |
| Brahman | 5 | 41.67% | 16.67% | 0.78% |
| Gelbvieh | 50 | 16.29% | 15.87% | 8.87% |
| Limousin | 38 | 100.00% | 15.02% | 21.11% |
| Jersey | 4 | 57.14% | 16.00% | 44.44% |
| Gir | 5 | 62.50% | 33.33% | 45.45% |
| Romagnola | 4 | 50.00% | 26.67% | 50.00% |
| Ndama | 4 | 57.14% | 36.36% | 50.00% |

**Table 2.** Variant density for assays used in this analysis before and after filtering.

| **Assay** | **Starting Assay Density** | **Filtered Density** |
| --- | --- | --- |
| GGP-ULD | 8,672 | 6,394 |
| GGP-LDv3 | 26,504 | 16,854 |
| SNP50 | 58,336 | 44,366 |
| GGP-90KT | 76,999 | 70,581 |
| GGP-HD | 139,977 | 125,446 |
| GGP-F250 | 227,234 | 201,236 |
| HD | 777,962 | 753,715 |

### Phasing and Imputation

#### Building imputation reference panels

After removing 308 individuals for testing, the remaining 28,183 F250 and 9,629 HD reference individuals (**Table 3**) were merged in PLINK, then phased with Eagle 2.4 using [27]. Missing genotypes inferred by Eagle were removed with bcftools [28] such that only phased, directly genotyped markers remained. The resulting phased F250 and HD genotypes were used as the imputation reference for “two-round” imputation, described below. The reference panel for “one-round” imputation was created in a recursive process by imputing missing HD markers for F250 assays, and missing F250 markers for HD assays with Minimac3, using all available observed HD and F250 assays as references respectively (**Supplementary Figure 1**).

**Table 3.** Breed representation of composite reference panel after removing 308 individuals for testing.

| **Breed** | **Complete**<br>**in CR** | **HD in**<br>**CR** | **F250 in**<br>**CR** | **% CR's**<br>**Complete** | **% CR's**<br>**HD** | **% CR's**<br>**F250** |
| --- | --- | --- | --- | --- | --- | --- |
| Holstein | 1932 | 3170 | 1944 | 80.13% | 32.92% | 6.90% |
| Gelbvieh | 257 | 265 | 514 | 10.66% | 2.75% | 1.82% |
| Angus | 132 | 2067 | 14454 | 5.47% | 21.47% | 51.29% |
| Simmental | 67 | 427 | 1759 | 2.78% | 4.43% | 6.24% |
| Brahman | 7 | 25 | 632 | 0.29% | 0.26% | 2.24% |
| Romagnola | 4 | 11 | 4 | 0.17% | 0.11% | 0.01% |
| Nelore | 3 | 855 | 4 | 0.12% | 8.88% | 0.01% |
| Jersey | 3 | 21 | 5 | 0.12% | 0.22% | 0.02% |
| Gir | 3 | 10 | 6 | 0.12% | 0.10% | 0.02% |
| Ndama | 3 | 7 | 4 | 0.12% | 0.07% | 0.01% |
| Brangus | 0 | 990 | 1603 | 0.00% | 10.28% | 5.69% |
| Hereford | 0 | 569 | 1834 | 0.00% | 5.91% | 6.51% |
| Mixed/crossbred | 0 | 419 | 2830 | 0.00% | 4.35% | 10.04% |
| Red Angus | 0 | 253 | 1905 | 0.00% | 2.63% | 6.76% |
| Limousin | 0 | 215 | 142 | 0.00% | 2.23% | 0.50% |
| Shorthorn | 0 | 136 | 218 | 0.00% | 1.41% | 0.77% |
| Charolais | 0 | 125 | 284 | 0.00% | 1.30% | 1.01% |
| Santa Gertrudis | 0 | 23 | 11 | 0.00% | 0.24% | 0.04% |
| Japanese Black | 0 | 19 | 0 | 0.00% | 0.20% | 0.00% |
| Brown Swiss | 0 | 15 | 0 | 0.00% | 0.16% | 0.00% |
| Norwegian Red | 0 | 5 | 0 | 0.00% | 0.05% | 0.00% |
| Chianina | 0 | 2 | 1 | 0.00% | 0.02% | 0.00% |
| Piedmontese | 0 | 0 | 9 | 0.00% | 0.00% | 0.03% |
| Braunvieh | 0 | 0 | 7 | 0.00% | 0.00% | 0.02% |
| Guernsey | 0 | 0 | 7 | 0.00% | 0.00% | 0.02% |
| Beefmaster | 0 | 0 | 3 | 0.00% | 0.00% | 0.01% |
| Sheko | 0 | 0 | 2 | 0.00% | 0.00% | 0.01% |
| Maine Anjou | 0 | 0 | 1 | 0.00% | 0.00% | 0.00% |

No phasing reference was used for within-breed phasing and imputation. The within-breed imputation reference consisted of 265 and 514 Gelbvieh individuals genotyped on the HD and F250 respectively. These reference assays, like above were merged, phased, and inferred genotypes were removed. A reciprocal F250/HD imputation with Minimac3 filled in missing genotypes in the reference.

#### Phasing and imputation

Reference-based phasing was performed for 308 downsampled individuals in Eagle using 9,629 pre-phased HD assays as reference haplotypes. To perform one-round imputation, phased assays were imputed against the complete, partially imputed 850K SNP composite reference using Minimac3 [29]. This imputation resulted in a total of 835,947 biallelic variants.

For two-round imputation, two separate rounds of imputation were performed to arrive at the 850K SNP density. First, downsampled and phased assays were imputed to HD density (759,329 SNPs), followed by a second round of imputation that inferred markers present on the F250, but not on the HD (122,181). The final imputed data (referred to hereafter as 850K) contained a total of 835,947 biallelic variants.

For within-breed imputation, 50 Gelbvieh animals with both F250 and HD genotypes, all of which were present in the multi-breed testing set, were downsampled to SNP50 density. They were phased with 1,113 additional Gelbvieh genotyped on the SNP50 using Eagle. This is representative of typical phasing strategies where a large number of individuals are genotyped at lower density. Phased genotypes were imputed against the breed-specific Gelbvieh reference (BR).

### Imputation Accuracy Calculations

Accuracy was measured in multiple ways for both individuals and variants for each imputation scenario. By coding alternate allele counts as 0, 1, and 2 (for AA, AB, and BB genotypes respectively), both correlations (R^2^) and count-based metrics could be used to evaluate the imputation accuracy of each variant and individual tested.

While simple concordance (ie. “correct/incorrect”) measures of accuracy are valuable, they overestimate the quality of imputation at low MAF and are ambiguous as to the nature of the error that creates an incorrectly imputed genotype. Instead of concordance, an imputation quality score (IQS) was calculated for each variant [30]. The IQS calculates concordances that are adjusted for the chance in guessing an imputed genotype correctly. This statistic provides similar conclusions to correlation for the majority of markers, but more robustly estimates imputation quality at low MAFs.

In addition to IQS, the exact nature of each error was cataloged and tallied for each individual and each variant. This allowed the errors to be categorized as either false heterozygotes (AA or BB imputed as AB), false homozygotes (AB imputed as AA or BB) or completely discordant (BB imputed as AA or vice versa). These more detailed error descriptions, in conjunction with minor allele frequency, position, and chip-of-origin information, allow for detailed analysis of how certain factors improve or reduce imputation accuracy to 850K in the context of each scenario.

To approximate how well represented an individual was in the composite reference, we created a centered genomic relationship matrix (GRM) as described in [31] using GEMMA [32]. We extracted diagonal elements of this matrix for each testing individual. The resulting values were quantitative measures of how diverged an individual was from the composite reference. Smaller values indicate individuals that are more closely related to the reference.

## Results

The minor allele frequency (MAF) spectrum of the HD and F250 assays that compose our reference panel is shown in **Figure 1** with the SNP50 assay for comparison. The SNP50 and HD have similar MAF spectra with mostly common variants. The HD assay also has an increased density of variants in the 0.025-0.075 MAF range. However, the F250 has a significantly higher proportion of its content with a MAF < 0.1, more similar to the site frequency spectrum of variants identified from genome resequencing.

### Imputation Accuracy Metrics

Numerous statistics are used to evaluate imputation quality. We compared two widely-used statistics (concordance rate and Pearson R^2^) with imputation quality score (IQS), a metric that has been used in multiple human studies, but not in livestock [30,33]. We tested each of these metrics on the TAUR dataset at the level of both variants and individuals. For variants, IQS was much more conservative than concordance, particularly at lower MAF (**Figure 2A**). The IQS was also more conservative than R^2^ across the MAF spectrum (**Figure 2B**). In the TAUR dataset, IQS scores were lower than their corresponding R^2^ values 81% of the time. At moderate to high MAF, these metrics mostly agree with one another, however when MAF < 0.1, both correlation and IQS penalize more heavily for imputation errors of rare variants, resulting in lower averages and higher variances in these two metrics compared with those observed for concordance. In addition to capturing these metrics, we also identify the type of error (complete discordance, false heterozygote, or false homozygote) that occurred.

**Figure 2.**
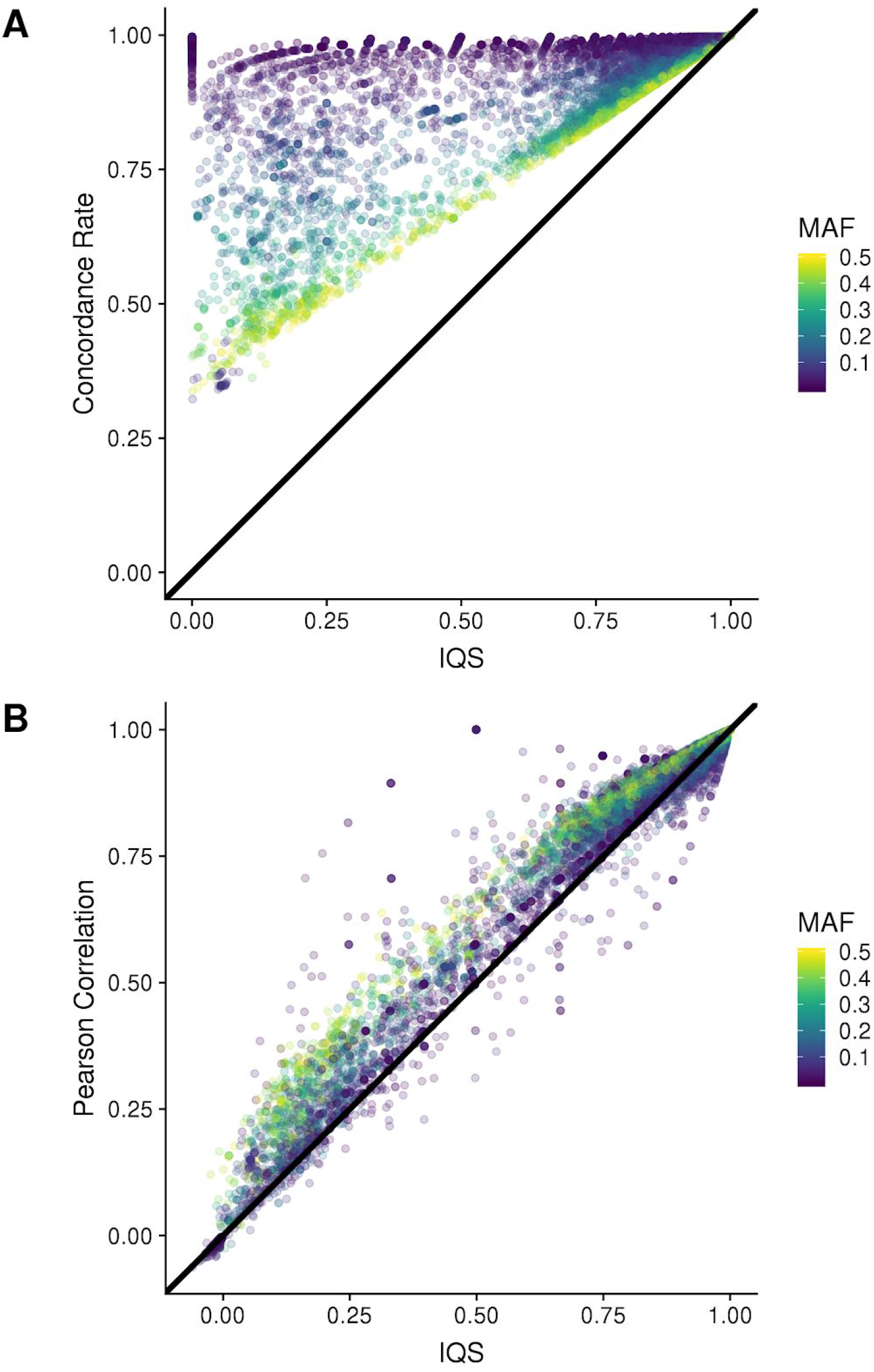
The IQS compared to concordance and correlation as measures of imputation accuracy. Three imputation accuracy measures calculated for the TAUR dataset. (A) Concordance and (B) Pearson correlation over-estimate imputation accuracies relative to the imputation quality statistic (IQS) resulting in bias and thus a false sense of high imputation accuracy.

Since IQS calculations do not work for assessing accuracy on the level of an individual, we used error type/count and Pearson R^2^ between observed and imputed genotypes to determine the impacts of different intrinsic and extrinsic factors on how well an individual is imputed. For our 308 testing animals, individual correlations ranged from 0.7466 to 0.9993. However, the majority of individuals had correlations of > 0.990. Individuals with the lowest R^2^ values (< 0.80) tended to have significantly more false heterozygote errors than false homozygote errors.

### Comparing Multi-Breed and Within-Breed Imputation Reference Panels

We used 49 Gelbvieh animals with observed HD and F250 genotypes that were downsampled to SNP50 genotypes to compare the accuracy of imputation between a multi-breed composite reference (CR) and a single-breed reference (BR) panel when imputing to 850K SNPs. SNP imputation accuracies were significantly higher for the CR compared to the BR panel. The breed-specific imputation panel performed quite well overall, with mean IQS score of 0.982. As a result, overall mean accuracy gains were modest but significant when using the CR (IQS=0.990) (**Figure 3A**). Of the 107,110 SNPs whose IQS changed when imputed against the different reference panels, 89,930 saw an increase with the CR, while only 15,349 were decreased. The average magnitude of these accuracy increases was significantly larger for the CR than for the BR (0.0797 vs. 0.0603).

**Figure 3.**
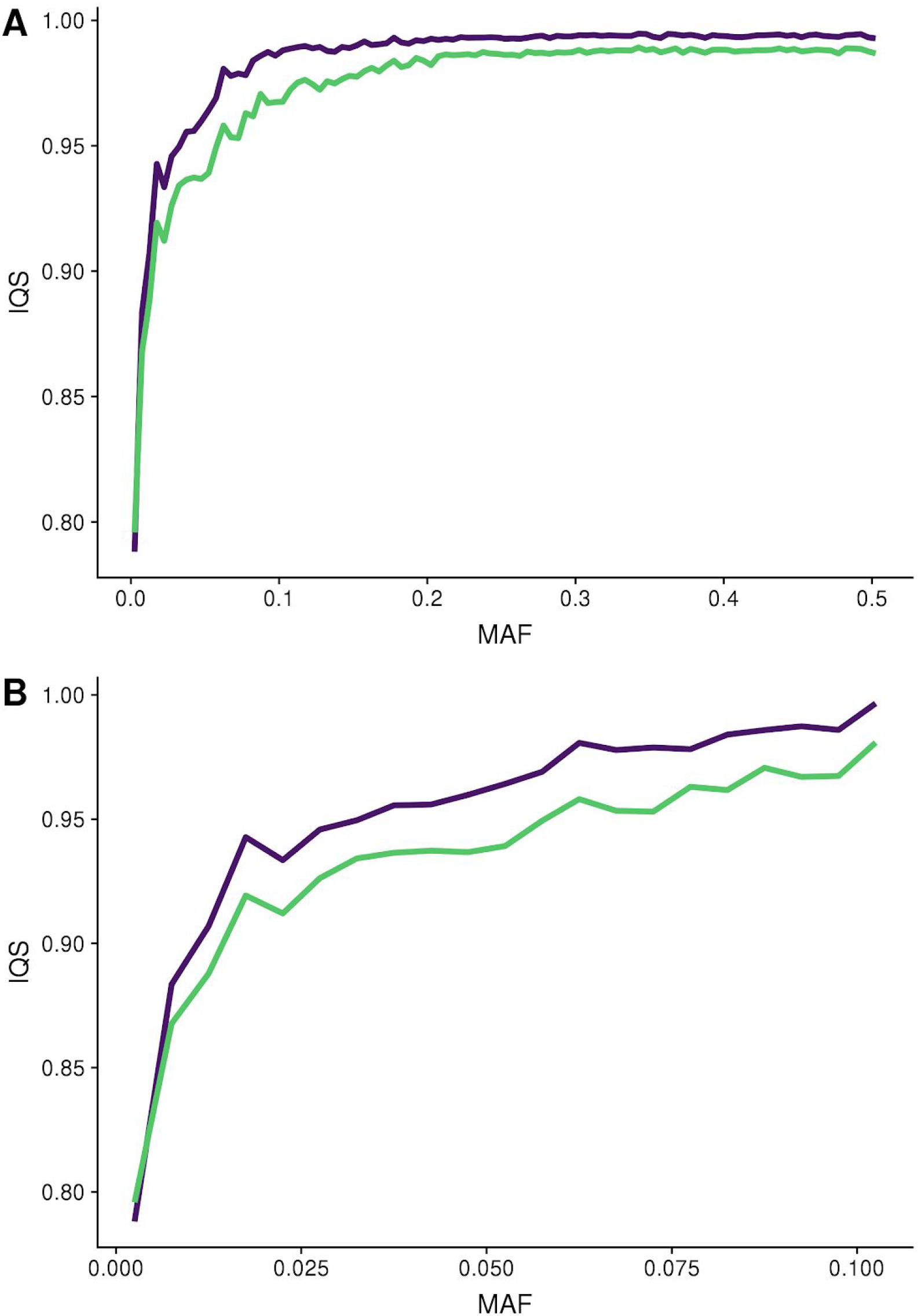
The composite reference improves per-variant imputation accuracies, particularly for rare variants. Imputation quality statistics when using breed-specific (green) and composite (purple) references for 850K imputation in the GEL dataset across the entire MAF spectrum (A), and at low MAF (B).

The most substantial accuracy gains from the CR were observed for low MAF variants (**Figure 3B**). While accuracy gains were modest for variants with MAF > 0.1 (0.007 IQS increase), rare variants saw a 0.0182 increase in IQS when imputing with a CR. This increase in the quality of low MAF imputation was not observed when using concordance or R^2^ statistics (**Table 4**). Of the 122,288 markers that were not perfectly imputed using the BR, there was an increase in IQS of 0.059 (R^2^ increase 0.032) when imputed with the CR.

**Table 4.** Per-variant imputation accuracy measures by MAF bin for GEL individuals imputed against BR and CR.

| MAF bin | BR | CR | BR R<sup>2</sup> | CR R<sup>2</sup> | BR IQS | CR IQS |
| --- | --- | --- | --- | --- | --- | --- |
|  | Concord. | Concord. |  |  |  |  |
| 0.00 - 0.05 | 0.999 | 0.999 | 0.984 | 0.982 | 0.910 | 0.926 |
| 0.05 - 0.10 | 0.997 | 0.998 | 0.985 | 0.99 | 0.959 | 0.979 |
| 0.10 - 0.15 | 0.996 | 0.998 | 0.986 | 0.992 | 0.974 | 0.989 |
| 0.15 - 0.20 | 0.995 | 0.997 | 0.988 | 0.993 | 0.982 | 0.991 |
| 0.20 - 0.25 | 0.994 | 0.997 | 0.990 | 0.995 | 0.986 | 0.993 |
| 0.25 - 0.30 | 0.994 | 0.997 | 0.990 | 0.995 | 0.987 | 0.993 |
| 0.30 - 0.35 | 0.994 | 0.997 | 0.991 | 0.996 | 0.988 | 0.994 |
| 0.35 - 0.40 | 0.993 | 0.997 | 0.992 | 0.996 | 0.988 | 0.994 |
| 0.40 - 0.45 | 0.993 | 0.996 | 0.992 | 0.996 | 0.988 | 0.994 |
| 0.45 - 0.50 | 0.993 | 0.996 | 0.992 | 0.996 | 0.988 | 0.994 |

One concern with using a large multi-breed reference panel for imputation is that it may introduce variation that does not actually exist in the population being imputed. To test this, we identify errors at the variant and individual levels as being either false heterozygotes or false homozygotes. Individuals had significantly fewer false heterozygote errors when using the CR compared to the BR (paired t-test p = 0.0039). There were on average 733 less false heterozygote calls per individual when using the CR.

While the per-variant imputation accuracy increases were significant, the most substantial improvements to imputation due to the CR were at the level of particular individuals. The mean individual correlations significantly increased from 0.9962 with the BR to 0.9979 using the CR (p = 0.0012). Animals that were already well-imputed using the BR did not show significant increases in accuracy with the CR. However, animals with the most BR imputation errors had much greater increases in accuracy with the CR (**Figure 4**). Of the 14 individuals who had > 5,000 total errors when using the BR, they had on average 5,522 fewer imputation errors with the CR indicating that these 14 individuals were significantly better imputed using the CR. Conversely, the 35 individuals with < 5,000 errors using the BR had only 209 fewer imputation errors on average when using the CR, indicating that the CR still performed better but to a lesser extent.

**Figure 4.**
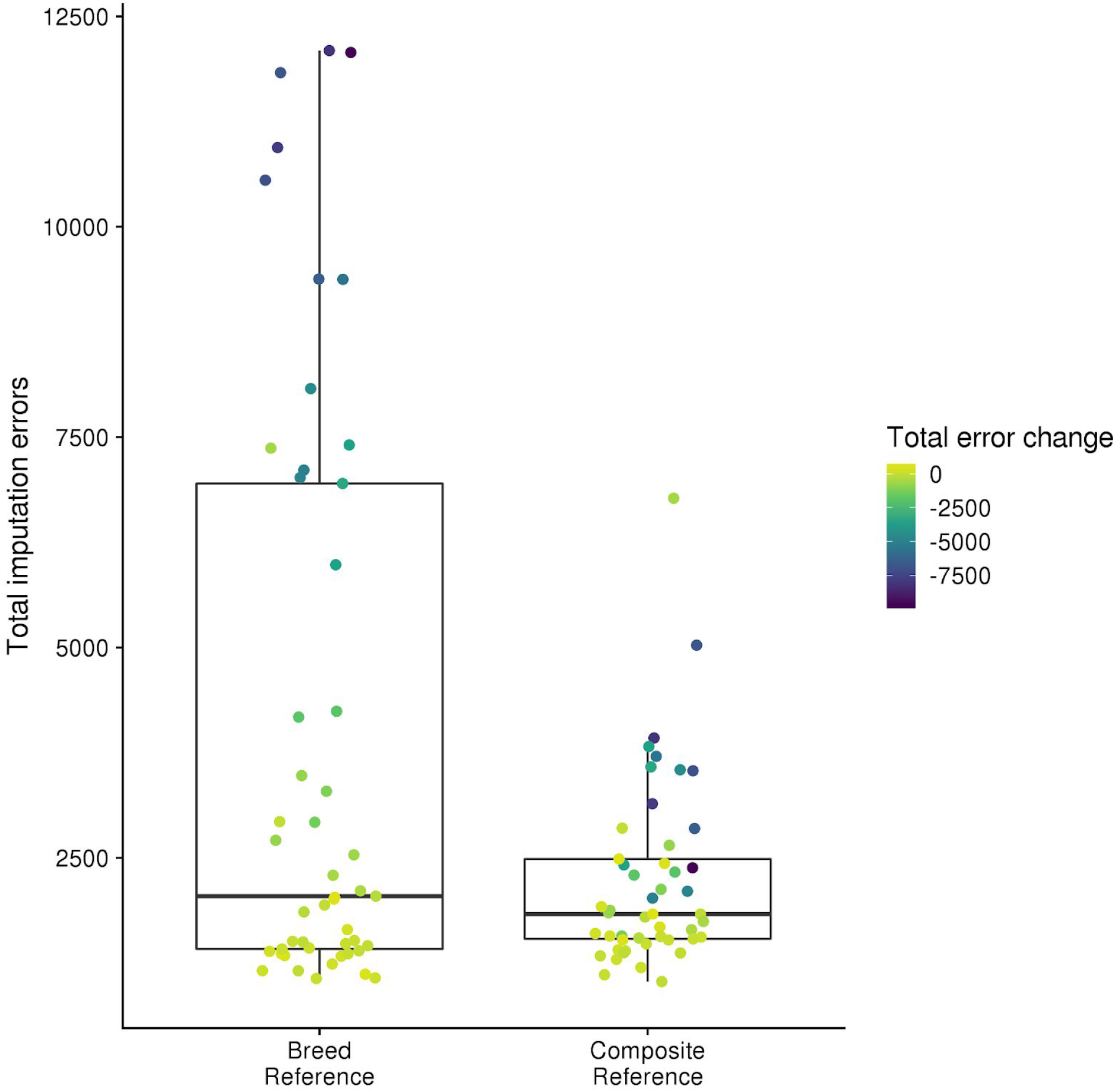
Improvements from the composite reference are largest for poorly imputed individuals. Comparing the total number of errors when imputing from SNP50 to 850K in GEL dataset when using breed-specific vs. composite references. Points are individuals, colored by the change in count of errors from breed to composite reference.

### One- vs. two-round imputation

To test the impact of utilizing partially imputed reference panels, we performed imputation on the TAUR testset of animals that were downsampled to SNP50 density using two different strategies for reference-creation (**Supplementary Figure 1**). Both strategies utilized the same set of HD and F250 reference genotypes. In our “one-round” imputation, the reference genotypes contain not only observed genotypes but also imputed genotypes. In our “two-round” imputation, samples are first imputed against a reference panel composed of 8,136 individuals with only observed HD genotypes, followed by a second round of imputation using 27,895 individuals with only observed F250 genotypes as a second reference. Both methods of imputation resulted in 835,947 total markers, 791,581 of which were imputed.

While the two-round imputation performed quite well, the one-round imputation outperformed the two-round imputation. Across the MAF spectrum, accuracies for the one-round imputation were consistently higher than those for the two-round method. However, the magnitude of these differences was modest. The one-round imputation increased the overall accuracy of imputation by 0.000762 IQS units, and the IQS at low MAF variants by 0.00256 units. The addition of rare variation from the F250 also increased the imputation accuracy in rare variants appearing exclusively on the HD.

The HD markers with MAF < 0.05 that were imperfectly imputed using the two-round method saw an average increase in IQS of 0.04248 (**Table 5**). For HD variants with moderate to high MAF, imputation accuracy is slightly increased using one-round compared to two-round.

**Table 5.** The IQS by MAF bin for HD-specific markers that are imperfectly imputed using two-round method.

| **MAF Bin** | **Number of SNPs** | **Two-round IQS** | **One-round IQS** | **IQS Change** |
| --- | --- | --- | --- | --- |
| 0.00 - 0.05 | 11,788 | 0.5453 | 0.6299 | 0.0425 |
| 0.05 - 0.10 | 18,095 | 0.9221 | 0.9330 | 0.0067 |
| 0.10 - 0.15 | 23,311 | 0.9724 | 0.9742 | 0.0017 |
| 0.15 - 0.20 | 29,222 | 0.9772 | 0.9784 | 0.0012 |
| 0.20 - 0.25 | 34,062 | 0.9803 | 0.9812 | 0.0009 |
| 0.25 - 0.30 | 39,948 | 0.9807 | 0.9815 | 0.0009 |
| 0.30 - 0.35 | 44,309 | 0.9815 | 0.9824 | 0.0009 |
| 0.35 - 0.40 | 47,657 | 0.9814 | 0.9822 | 0.0008 |
| 0.40 - 0.45 | 49,233 | 0.9815 | 0.9823 | 0.0008 |
| 0.45 - 0.50 | 50,908 | 0.9816 | 0.9823 | 0.0007 |

### Impact of breed representation in reference panel on imputation accuracy

Using individual imputation accuracy measures for 308 test animals, we identify the effects of an individual’s breed composition and those breeds’ representations in the CR on how well an individual is imputed. Using the CR, individual R^2^ ranged from 0.747 to 0.999 while total imputation errors ranged from 932 to 219,737. The accuracy of imputation appeared to be strongly related to an animal’s labeled breed (**Table 6**). Individuals from breeds adequately represented in the CR (Angus, Gelbvieh, Hereford, Holstein, Jersey, Limousin, Nelore, and Simmental, **Table 3**) are generally well imputed (median R^2^ = 0.997, range = [0.930, 0.999]) (**Figure 5**). Gelbvieh individuals had the highest mean imputation accuracy of any breed (R^2^ = 0.998). This is likely a function of the high proportion of Gelbvieh animals genotyped on both the F250 and HD in our reference panel. Gelbvieh made up 10.66% of our reference animals with complete 850K genotypes, second only to Holstein (80.13% of total). Since HD markers make up the largest proportion of the 850K SNP set, individuals from breeds with many HD genotypes, but few F250 genotypes, like Nelore, were still imputed at high accuracy (R^2^ = 0.981). Individuals from breeds that are sparsely represented in the CR (Brahman, Gir, N’Dama, and Romagnola) show decreases in means and increases in the variance of per-animal imputation accuracies (mean R^2^ = 0.890, range = [0.747, 0.961]).

**Table 6.** Composite reference 850K imputation, individual accuracies (R^2^) by breed.

| **Breed** | **Mean**<br>**Correlation** | **Min**<br>**Correlation** | **Max**<br>**Correlation** |
| --- | --- | --- | --- |
| Gelbvieh | 0.9979 | 0.9935 | 0.9989 |
| Hereford | 0.9971 | 0.9912 | 0.9988 |
| Holstein | 0.9969 | 0.9947 | 0.9984 |
| Simmental | 0.9963 | 0.9841 | 0.999 |
| Angus | 0.9953 | 0.959 | 0.9993 |
| Jersey | 0.995 | 0.9905 | 0.9966 |
| Limousin | 0.9892 | 0.93 | 0.996 |
| Nelore | 0.981 | 0.9774 | 0.9844 |
| Brahman | 0.9412 | 0.932 | 0.9611 |
| Gir | 0.9027 | 0.8689 | 0.9482 |
| Romagnola | 0.8742 | 0.8549 | 0.8958 |
| N'Dama | 0.7632 | 0.7466 | 0.8033 |

**Figure 5.**
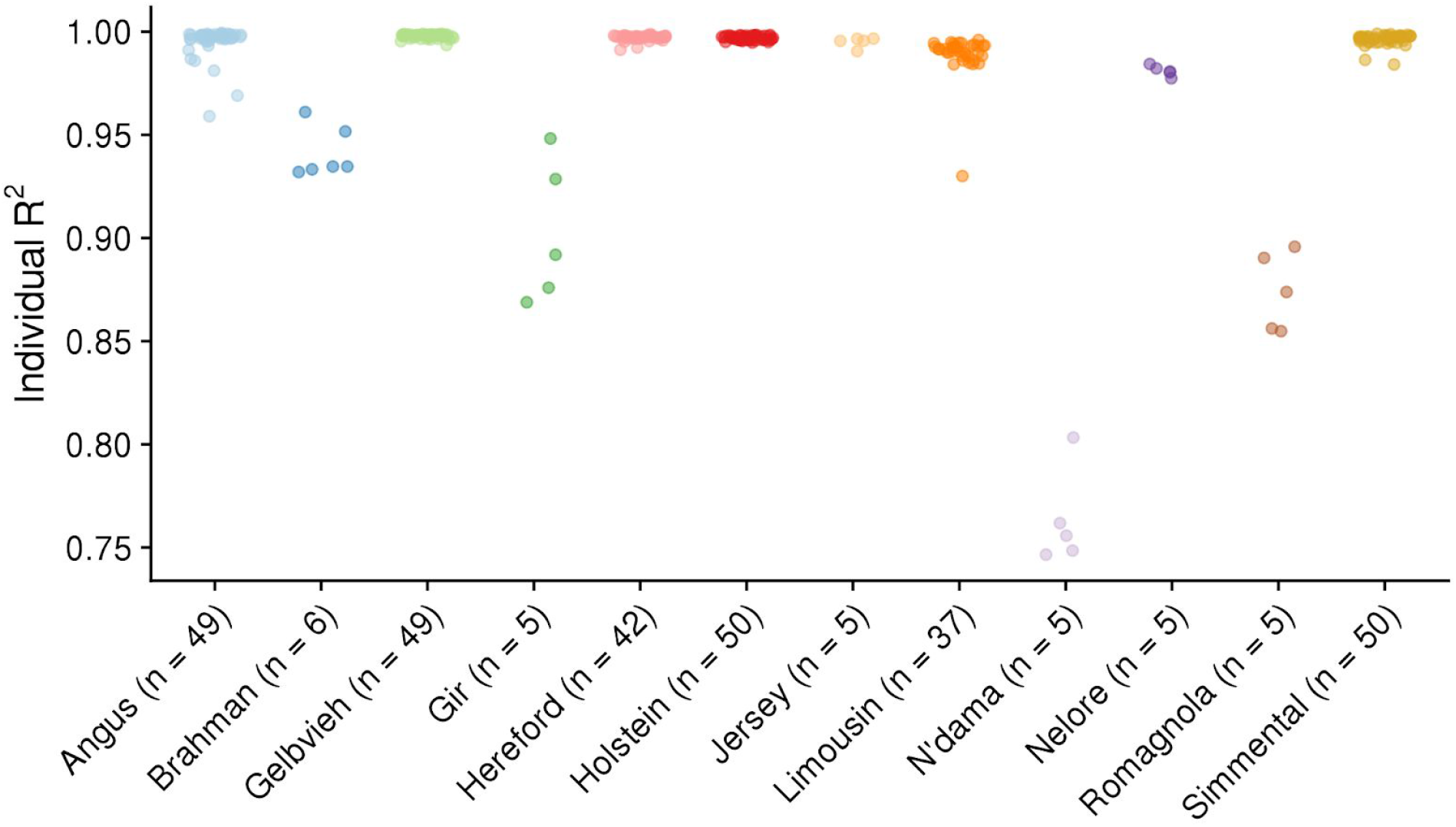
Per-individual accuracy by reported breed. Individual Pearson R^2^ by breed. Each point is an individual, colored by breed.

We used a genomic relationship matrix (GRM) created with observed genotypes from all reference and testing individuals to test if an individual’s increased relatedness to the CR would translate to increased imputation accuracies (**Figure 6**). There was no clear relationship between an individual’s overall relatedness to the CR and imputation accuracy. Instead, imputation accuracy was better predicted by the breed representation of the individual in the CR. For example, individuals assigned as Romagnola had relatively low imputation accuracies (mean individual R^2^ = 0.874), even though they were highly related to the CR. Their low imputation accuracy likely stems from a low representation of Romagnola haplotypes in the CR (15 HD and 8 F250 genotypes). The converse was seen in animals labeled Nelore. Although they were distantly related to the CR as a whole, the higher number of samples in the reference panel (858 HD and 7 F250 genotypes) resulted in accurate imputation overall (mean individual R^2^ = 0.981). This is further illustrated by Gir, which are similarly diverged as Nelore but have lower accuracy due to only 13 HD and 9 F250 individuals in the CR. Further, even low levels of admixture with breeds not adequately represented in the CR can lead to decreased imputation accuracies. Breed composition information was valuable for identifying outlying individuals or breeds that, in theory, should have been imputed accurately. For example, the five individuals labeled as Angus with outlying imputation accuracies can be explained by admixture with breeds not well represented in the CR (Supplementary Table 2). Each of the observed outliers labeled as Angus had relatively low proportions of Angus ancestry (0.0763 to 0.3169), and moderately high proportions of breeds sparsely represented in the CR.

**Figure 6.**
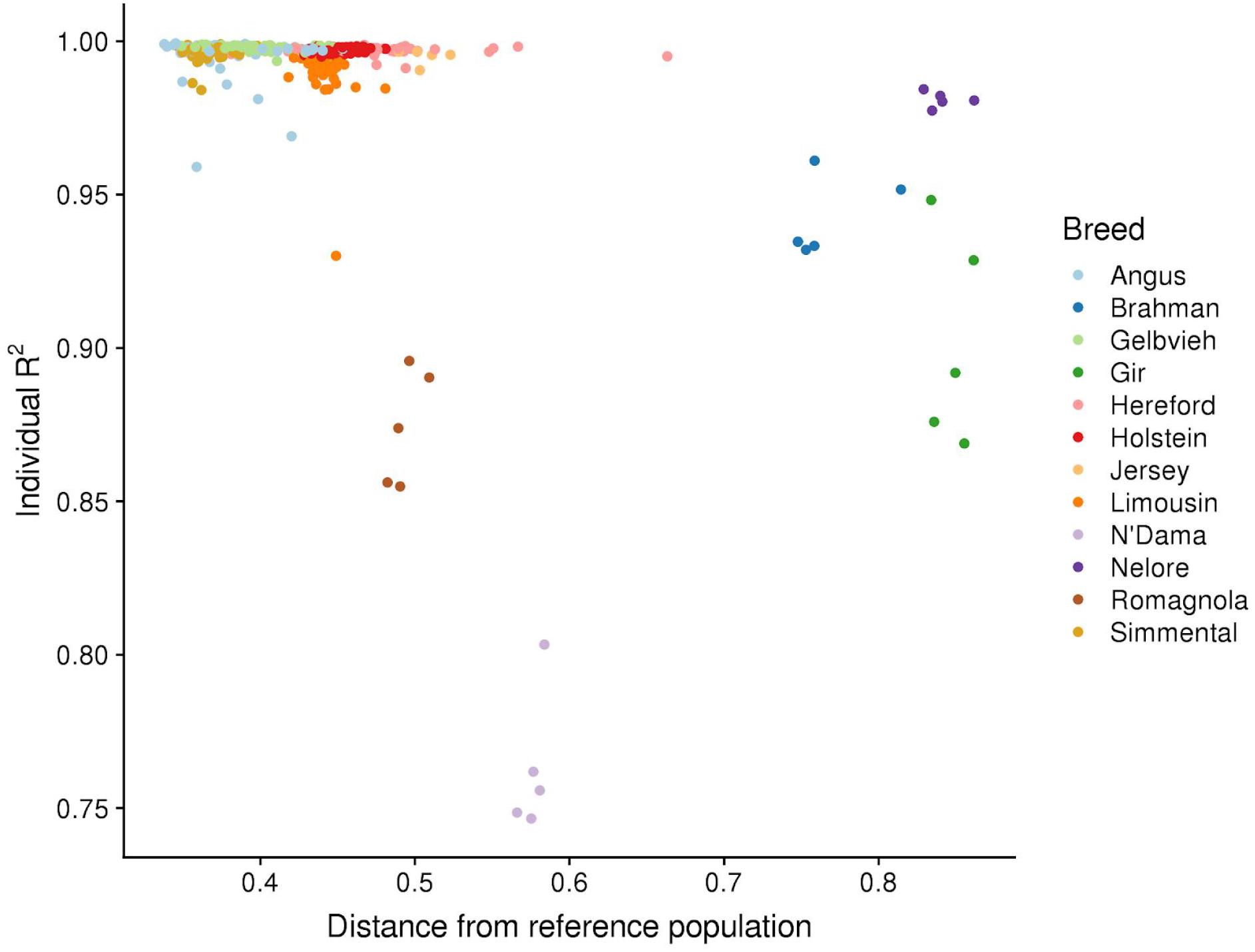
Relatedness to whole composite reference is not best predictor of individual imputation accuracy. By-individual R^2^ as a function of individual’s divergence from GRM calculated from composite reference.

### Impact of starting assay density on 850K imputation

The starting density of an assay affects the accuracy of imputation to HD density as well as sequence-level. To test the impact of starting chip density on 850K imputation accuracy, we used the TAUR dataset downsampled to 5 common commercial assay densities. Each successive increase in chip density yielded increases in imputation accuracy both overall and for low-MAF variants (**Table 7** and **Figure 7**). The largest increase in accuracy came between the ULD and GGPLD densities. Variant accuracies when imputing from the ULD were exceptionally poor at low minor allele frequencies. While the decline in IQS at low MAF was seen with other starting densities, it was much stronger with the ULD (0.1385 IQS decrease). The overall gains in imputation accuracy when moving to higher density assays (SNP50, GGP90KT, GGPHD) are minimal, but increases to low MAF marker imputation are more substantial.

**Table 7.** Per-variant mean and standard deviations for imputation quality statistics for 850K imputation in TAUR dataset based on starting assay density.

| **Starting Assay** | **Starting Density** | **Mean IQS** | **SD IQS** | **Mean IQS (Low MAF)** | **SD IQS (Low MAF)** |
| --- | --- | --- | --- | --- | --- |
| ULD | 6,394 | 0.9095 | 0.1766 | 0.7720 | 0.3503 |
| GGPLD | 16,854 | 0.9612 | 0.1225 | 0.8969 | 0.2604 |
| SNP50 | 44,366 | 0.9745 | 0.1154 | 0.9121 | 0.2468 |
| GGP90KT | 70,581 | 0.9796 | 0.1104 | 0.9220 | 0.2402 |
| GGPHD | 125,446 | 0.9832 | 0.1032 | 0.9319 | 0.2264 |

**Figure 7.**
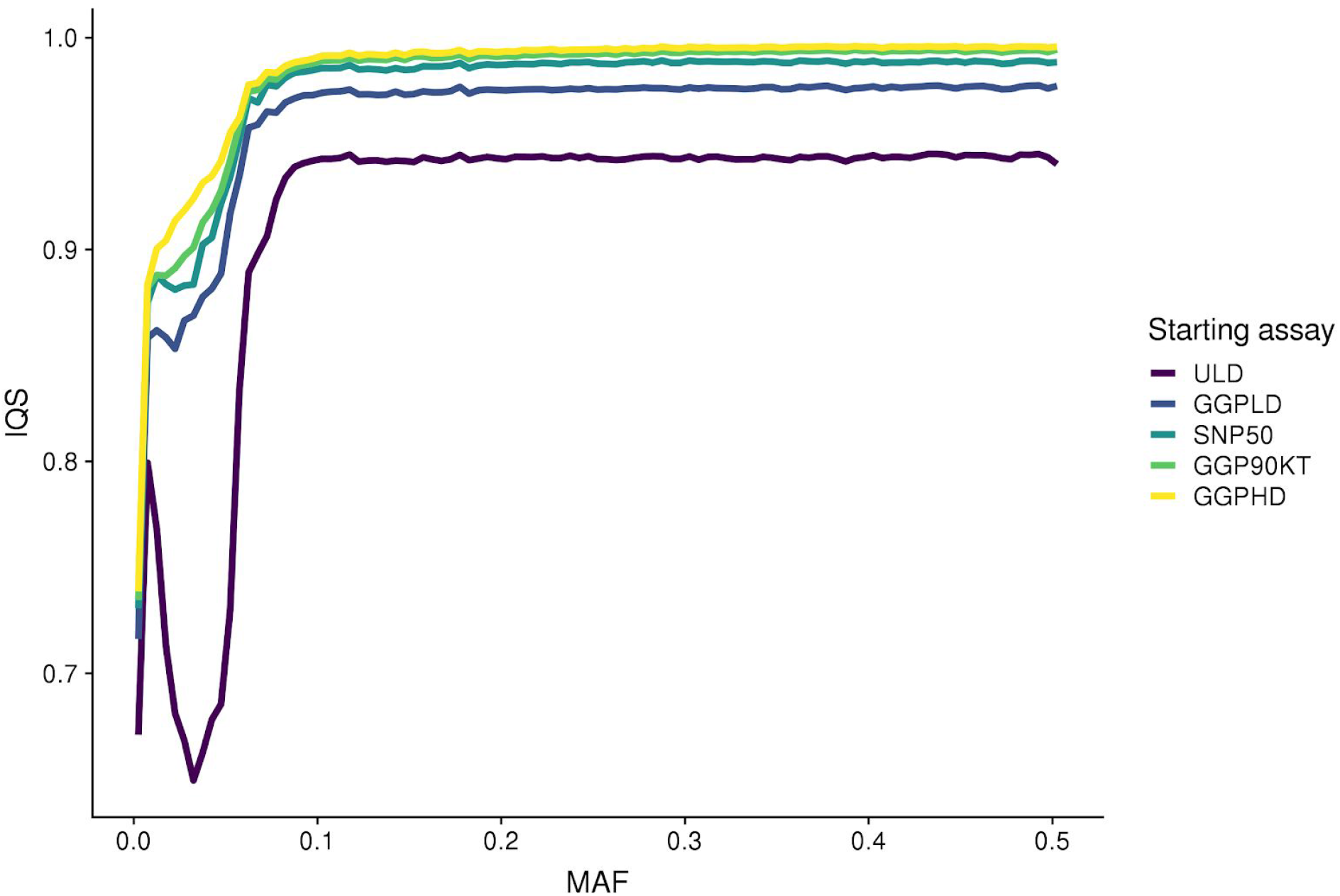
Effects of starting assay density on imputation accuracy across MAF spectrum. Variant accuracy measures for 850K imputation in TAUR dataset based on five assay starting densities. Binned mean IQS lines (per-variant accuracy) across MAF spectrum.

Individual accuracies also increased with assay starting density. A one-way ANOVA using Tukey’s method for multiple comparisons showed that there was a significant difference in 850K imputation accuracy between the ULD and GGPLD (p = 9.05e-5), but not between GGPLD and SNP50 (p = 0.1486). (**Figure 8**). No significant difference existed between the SNP50 and GGP90KT or GGPHD. The GGPLD starting density was, however, significantly different from the GGP90KT (p = 0.0049). This suggests, like above, that accuracy gains are minimal when the starting density is greater than 50,000 variants. Median individual R^2^ values are given in **Table 8**.

**Figure 8.**
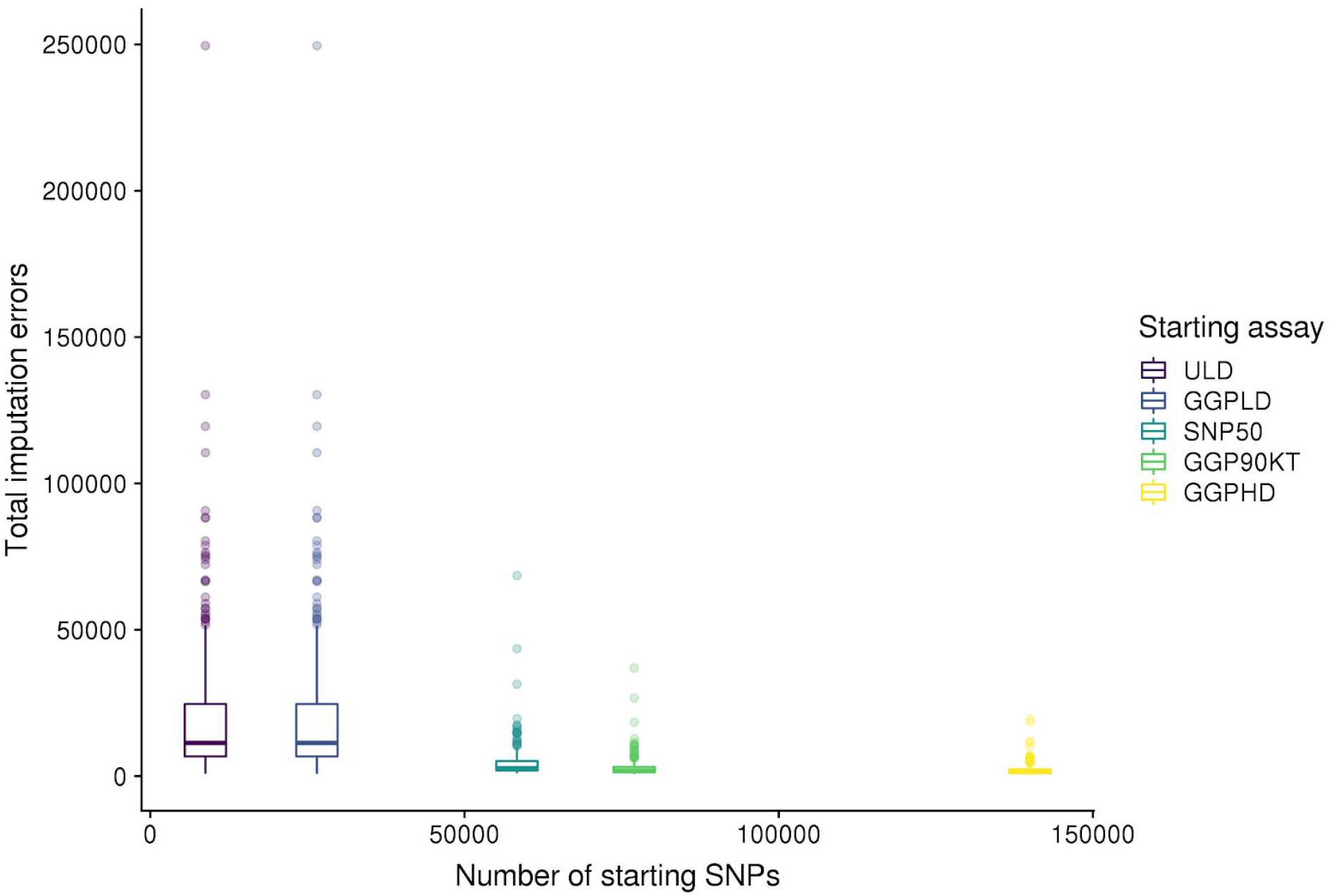
Impact of starting density on per-individual imputation accuracy. Per-individual accuracy measures for 850K imputation in TAUR dataset based on five starting assay densities. Boxplots for total imputation errors based on each starting assay density.

**Table 8.** Per-individual R^2^ mean and standard deviation values for 850K imputation based on starting assay density.

| **Starting Assay** | **Starting Density** | **Median $R^2$** | **SD $R^2$** |
| --- | --- | --- | --- |
| ULD | 6,394 | 0.989 | 0.6070 |
| GGPLD | 16,854 | 0.995 | 0.0486 |
| SNP50 | 44,366 | 0.997 | 0.0314 |
| GGP90KT | 70,581 | 0.998 | 0.0205 |
| GGPHD | 125,446 | 0.999 | 0.0129 |

### Error profiles and regions of low imputation accuracy

Using the imputation accuracy information for the TAUR dataset, we identify a number of regions of low imputation accuracy and diagnose why they occur. While most markers are imputed correctly, most chromosomes have at least one small region of poorly imputed markers (**Figure 9**). The overall number of poorly imputed markers was quite low. Only 21,848 markers had IQS < 0.8 (1.95% of imputed makers), and only 8,963 had more than 10 total imputation errors (1.07% of imputed markers). When using the IQS metric, we see that low-accuracy markers exist across each chromosome, particularly low MAF variants with relatively few errors (making IQS = 0) (**Figure 9A**). However, both IQS and total errors (**Figure 9B**) show peaks of low imputation accuracy. Investigation of these regions indicated that the probe sequences for these variants had multiple equally likely matches to the genome, which is indicative of either genome mis-assembly or the wrong location was chosen to represent the placement of the marker. The latter can be easily rectified based on changing the map files for these variants to reflect the alternate placement.

**Figure 9.**
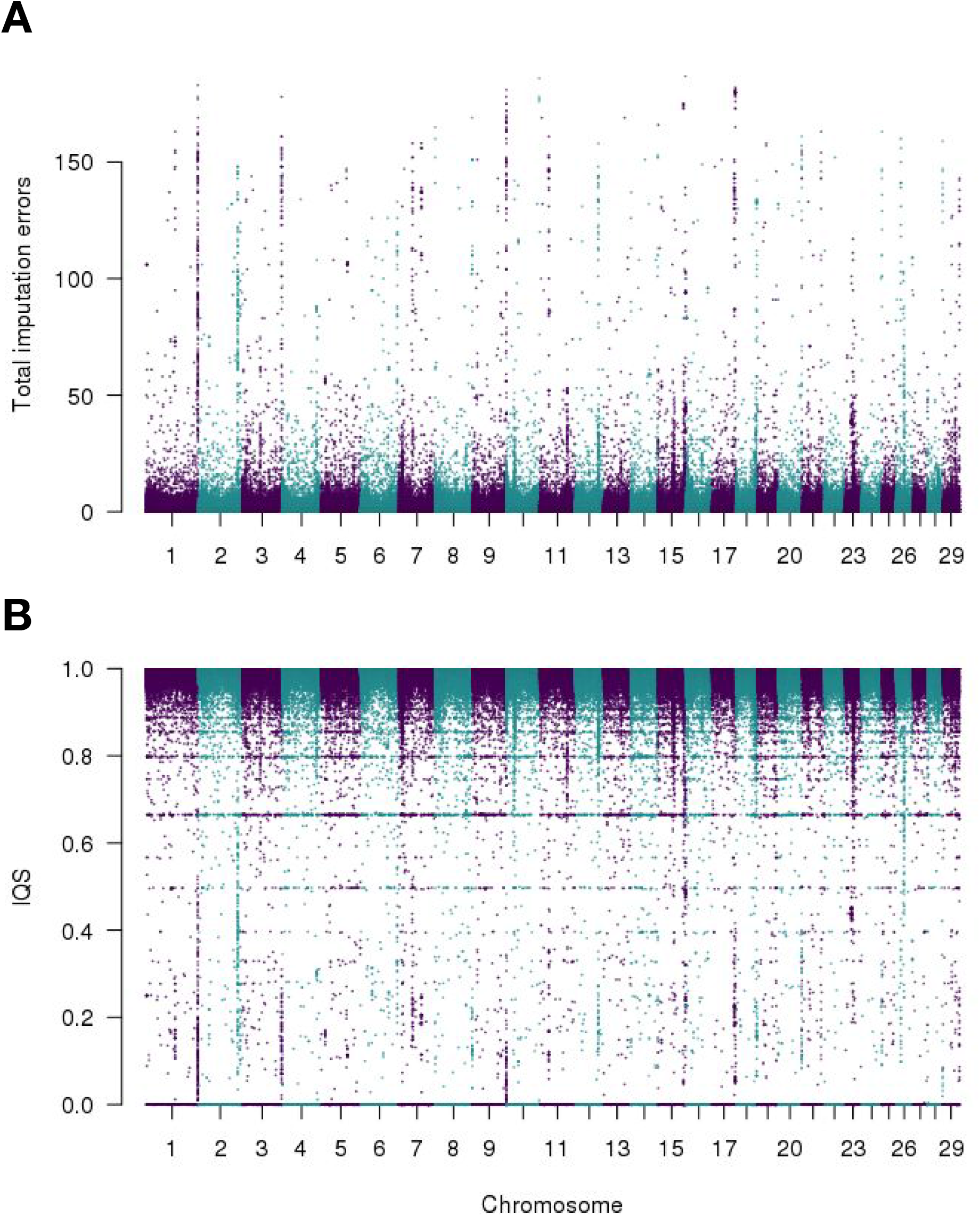
Regions of low imputation accuracy exist across the genome but are a small subset of markers. Regions of low imputation accuracy identified for the TAUR dataset using total imputation errors (A), and IQS (B).

## Discussion

### Imputation Accuracy Metrics

Most studies involving imputation accuracy in livestock populations use two methods for assessing the correctness of imputation: concordance, the proportion of correctly imputed genotypes, and Pearson correlation between observed and imputed genotypes. While both of these statistics make sense at the level of individuals, their ability to identify poorly imputed markers is suboptimal, particularly at low minor allele frequencies. Because of the nature of our data, which contains a large proportion of rare variants, a statistic that more robustly represents the actual quality of imputation is essential. Using the imputation quality statistic (IQS) [30], we show that Pearson R^2^ and especially concordance rate, overestimate the accuracy of imputation at low MAF variants. Unlike R^2^, which requires variation, IQS can be calculated for variants where no variation exists in either the observed or imputed dataset but does in the other. Of these, 2,070 variants existed in the GEL testing data. Their average MAF was 0.040, meaning that they had high concordances (0.97 average), but very low IQS scores (0.0). This information would be lost when using R^2^, and grossly inflated if using a concordance value. It is worth noting that for this work, we treat HD and F250 genotype calls as being correct, even though there is approximately a 0.5% error rate associated with these genotyping methods [12].

### F250 impact on rare variant imputation

The F250 was designed to assay a large number of rare, potentially functional variants and therefore is very gene-centric. Common variants were included on the F250 assay to allow for imputation and genomic prediction applications. The rare variants present on the F250 are important in the context of this work for two main reasons. First, imputing an additional ∼170,000 variants on a population level will increase researchers ability to refine GWAS signals and identify putative QTN due to increased marker density. Second, because the variants are very gene-centric, it is anticipated that imputation to whole genome sequence level will be improved within genic regions. Inclusion of rare variants may increase the imputation accuracy of other rare variants that are not directly assayed as linkage disequilibrium (r^2^) is maximized when allele frequencies between markers are similar. Rare alleles in the absence of selection are assumed to be more recent, and thus likely to be in linkage disequilibrium with other recently acquired rare alleles [34]. By adding rare variation to our reference panel with the F250 assay and genotyping a large number of individuals, we improve the imputation of rare variants that aren’t directly assayed by the F250. Even though many individuals in our reference panel only have imputed F250 genotypes, their presence has a significant impact on rare variant imputation accuracies. We expect to see this boost in rare variant imputation accuracy from the F250 carry over to subsequent imputation to sequence-level. The impact of F250 on rare variant imputation underlines the need for more complete 850K data in our reference (individuals genotyped on both the HD and F250). Imputation accuracies are the highest in breeds with the largest numbers of complete genotypes because more of the haplotypic diversity in those breeds is captured.

### Multi-breed vs. within-breed imputation reference panels

Early studies involving imputation concentrated mostly on homogenous populations. When using closely related animals from a population like Holstein [35,36], high-quality imputation can be achieved from a relatively small reference set of genotypes, provided those animals are closely related to the animals being imputed. In recent years, large numbers of low-density genotypes from many cattle populations, including outbred animals, as well as admixed individuals, both registered and commercial have been generated. Along with these data, many animals, from a wide range of breeds have been genotyped on high-density assays like the HD and F250. By combining all available high-density genotypes into a single multi-breed composite reference panel, we see increases in imputation accuracy across the MAF spectrum. The most substantial increases when using the CR compared to a breed-specific reference came on the level of specific individuals. Samples that were imputed well against the breed reference did not see substantial increases in accuracy when imputed against the CR. However, individuals poorly imputed using the BR saw a substantial reduction in the number of imputation errors when using the CR. The increased haplotypic diversity of the composite reference acts precisely as would be expected, by improving the imputation of haplotypes that would not have been found in a breed-specific reference. It is important to note that in the context of routine genotyping and imputation, one will not know *a priori* which individuals may have poorly imputed genotypes. Gains in accuracy when using the CR will likely be small, but never worse than a breed-specific reference, in purebred, closed herdbook populations like Holstein or Angus, but for open herdbook or composite breeds, we expect to see substantial increases. Further, we did not see an increase in false heterozygous or false homozygous genotypes when including multiple breeds in the reference panel, indicating that the CR is not introducing variation not present within the imputed individuals. For breeds adequately represented in the CR, we see imputation accuracies (median R^2^ = 0.997) approaching the error rate for genotyping assays (0.5%).

The success of our composite reference panel for 850K imputation also points toward strategies for sequence-level imputation. Previous work has advised the use of multi-breed reference panels for sequence imputation [37,38]. Our work at the high-density genotype level confirms this and suggests that improvements in imputation for outbred and admixed populations will benefit from the sequencing and inclusion of “non-core” animals that represent a greater amount of haplotypic diversity.

### Breed representation in reference

An individual’s relatedness to the entire CR was not a good predictor of imputation accuracy. Our multi-breed reference is heavily biased towards the most common and economically relevant American beef breeds but has a diverse array of other breeds in varying numbers. There appears to be a point at which an individual’s genomic representation in the reference reaches a threshold and meaningful gains in accuracy cease. At this point, the most significant gains in accuracy will likely come through the addition of high-density genotypes for breeds sparsely represented in our reference, or through the addition of more completely genotyped individuals, i.e. those with both HD and F250 genotypes. It is worth noting that the by-breed accuracies reported here for populations with limited representation in our composite reference (Brahman, Gir, N’Dama, Romagnola) would have likely performed better had we not removed large proportions to be a part of our testing set. Therefore, we expect these accuracies to be underestimates should individuals from these breeds be imputed using the full CR.

### Starting Assay Density

The starting density of an assay has a significant impact on the accuracy of imputation to 850K. In agreement with the Bovine HapMap project, we see that approximately 50,000 SNPs are needed to impute to higher densities with high accuracy [39]. This observation likely has a larger impact on research applications seeking to identify QTN, compared with applications for genetic prediction. There are a large number of individuals that have been genotyped at extremely low densities (< 10,000 markers) and it has been demonstrated that these individuals can be accurately imputed to ~50,000 markers [35,36]. For 850K and sequence imputation, we recommend that these individuals are first imputed to the level of ~50,000 variants within-breed to leverage the linkage information present within these large pedigrees, such as the Holstein breed and the resulting genotypes can be accurately imputed using the CR to the 850K level. However, groups wishing to perform both genomic predictions and downstream causal variant discovery, via imputation, would be best served genotyping new individuals with an assay density of ~50,000 SNPs.

## Conclusions

We conclude that in diverse samples, as seen in typical beef cattle populations, a multi-breed phasing and imputation panel will provide the highest accuracies. Individuals whose ancestry is moderately represented in the reference are imputed exceptionally well. By-variant imputation accuracies were highest for rare variants when using the composite reference. The addition of rare variation from the F250 assay further increased the accuracy of rare variant imputation from the HD assay. The addition of a large number of individuals genotyped for rare variants will likely have similar effects on rare variant imputation at the sequence level as well. We confirm that for 850K imputation, significant gains in accuracy plateau when increasing the starting-assay density past 50,000 SNPs. We have identified a small subset of SNPs that have poor imputation accuracies, most of which are caused by probe sequence location errors that will be fixed. Further improvement in accuracy can be obtained by removing Mendelian inconsistencies from the raw data used to create the CR, which was not done for this work. Furthermore, the largest gains in accuracy are expected to come from the addition of individuals with complete genotypes (HD and F250), with the largest realized gains from even a modest increase in the less well-represented breeds. Imputation accuracies for breeds adequately represented in the multi-breed composite-reference panel with a starting assay density of at least 50,000 SNPs should experience accuracies approaching the error rate of genotyping arrays. We anticipate the CR presented here can serve as a foundation reference panel with which the global cattle community can build upon to impute lower density genotypes in a consistent and accurate manner.

## List of Abbreviations

CR: = composite reference
BR: = breed reference
MAF: = minor allele frequency
GEL: = Gelbvieh testset
TAUR: = taurine testset
SNP: = single nucleotide polymorphism
IQS: = imputation quality statistic

## Declarations

### Ethics approval and consent to participate

Not applicable

### Consent for publication

Not applicable

## Availability of data and material

### Funding

This project was supported by Agriculture and Food Research Initiative Competitive Grant no. 2016-68004-24827 from the USDA National Institute of Food and Agriculture.

### Authors’ contributions

**Supplementary Table 1.** Shared markers between analyzed assays. Counts of shared, unfiltered markers between assays used in this analysis.

|  | **HD** | **F250** | **GGPHD** | **GGP90KT** | **SNP50** | **GGPLD** | **ULD** |
| --- | --- | --- | --- | --- | --- | --- | --- |
| **HD** | **777,962** | 37,841 | 134,599 | 74,524 | 50,665 | 25,693 | 8,280 |
| **F250** |  | **227,234** | 31,156 | 19,176 | 22,199 | 18,774 | 8,070 |
| **GGPHD** |  |  | **139,977** | 73,341 | 42,975 | 25,121 | 8,151 |
| **GGP90KT** |  |  |  | **76,999** | 28,952 | 14,240 | 7,948 |
| **SNP50** |  |  |  |  | **58,336** | 8,607 | 8,034 |
| **GGPLD** |  |  |  |  |  | **26,504** | 8,320 |
| **ULD** |  |  |  |  |  |  | **8,762** |

**Supplementary Table 2.**
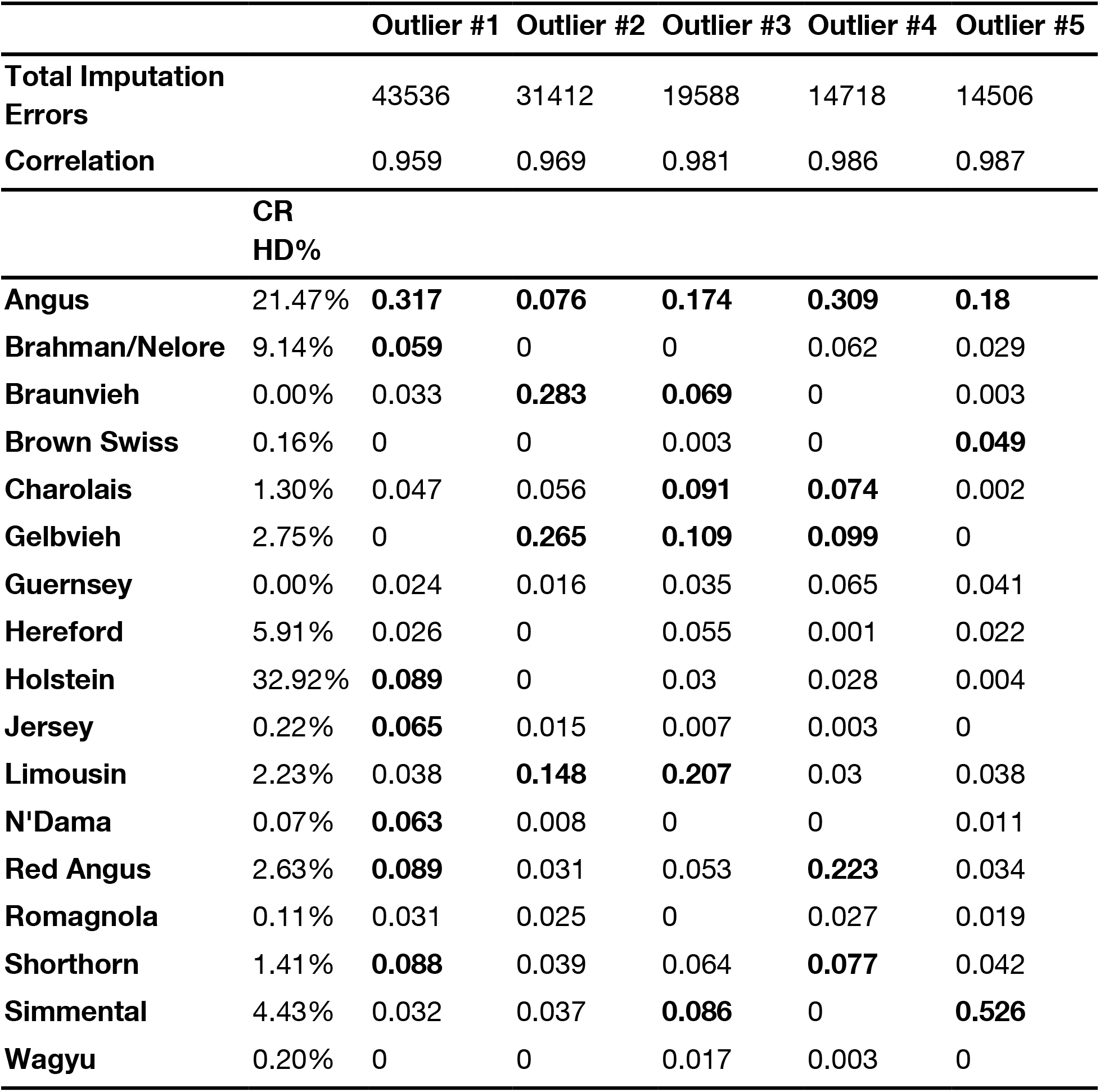
Outlier samples labelled Angus and their corresponding breed composition. Bolded values represent the largest values that sum to 75% of an individual’s total breed composition. The percentage of individuals from each breed with HD genotypes in the CR is indicated.

**Supplementary Figure 1.**
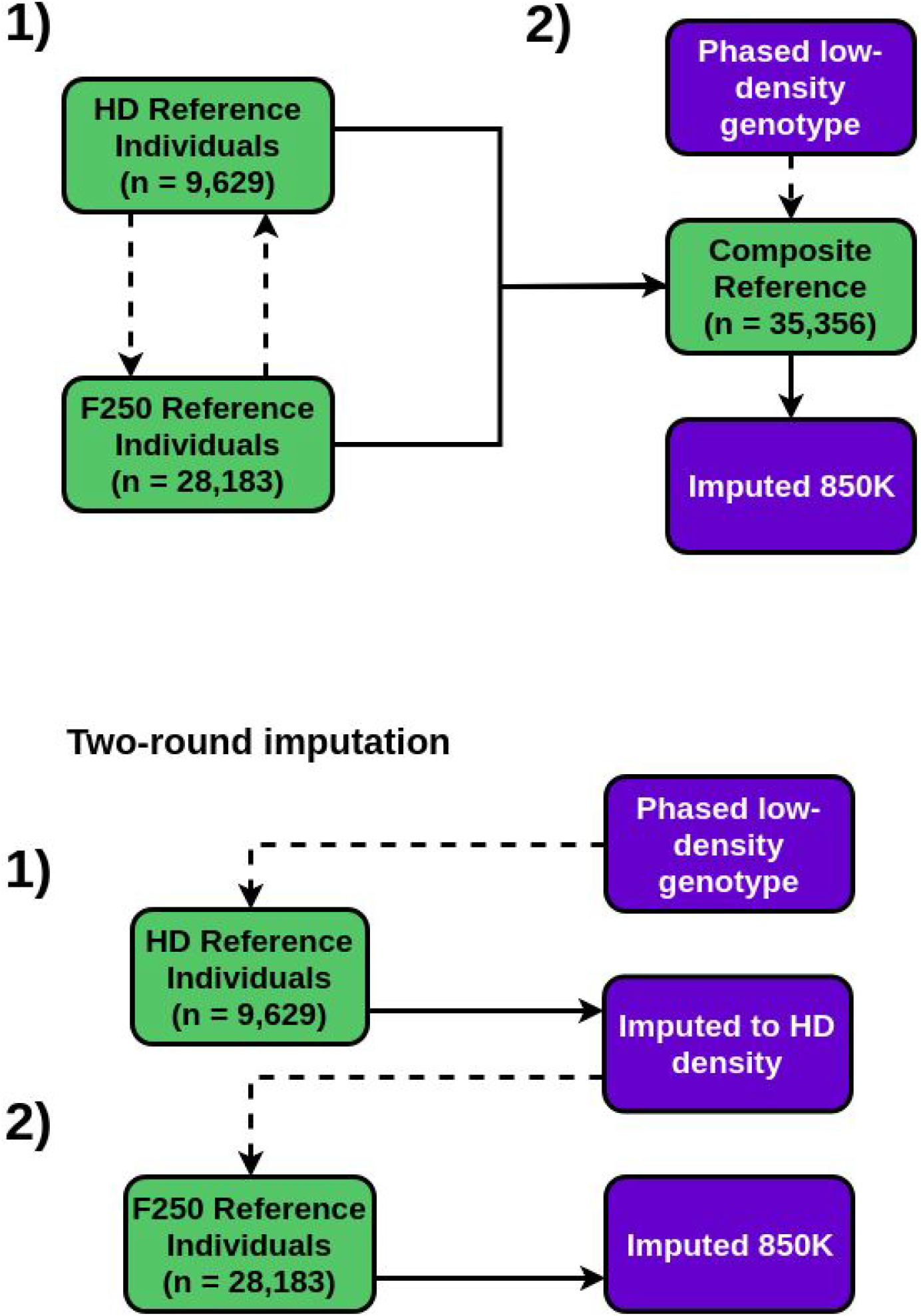
Schematic representation of one vs. two-round imputation. Dotted lines represent imputation. In one-round imputation (A), HD and F250 reference samples are cross-imputed to create a partially imputed composite reference (1). This is followed by a single round of imputation of low-density genotypes using the CR (2). For two-round imputation (B), two rounds of imputation occur: first from low-density to HD (1) and then from HD to 850K (2).

**Supplementary Figure 2.**
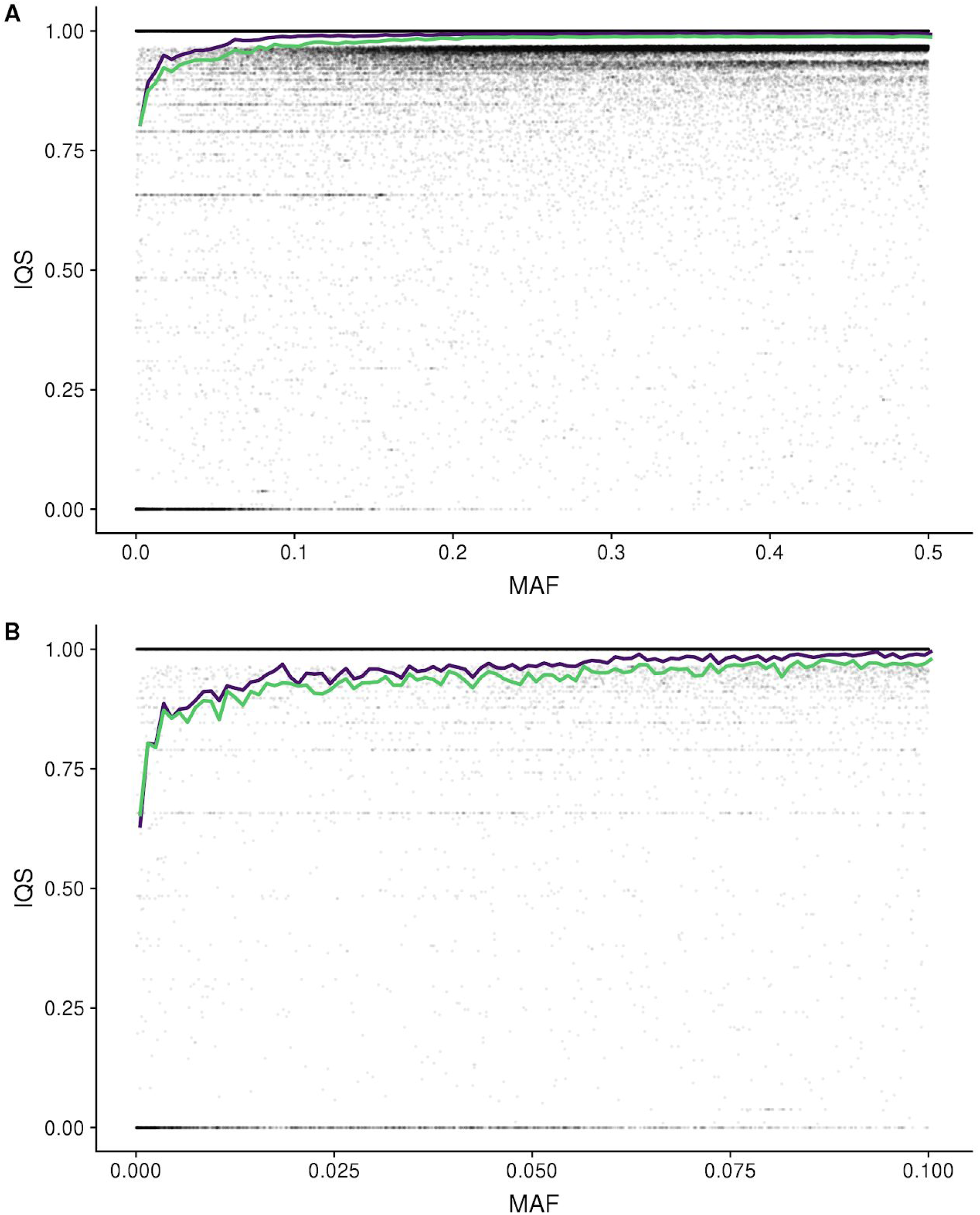
Imputation quality statistics when using breed-specific (green) and composite (purple) references for 850K imputation in the GEL dataset across the entire MAF spectrum (A), and at low MAF (B). Points are individual variants.

